# Ambient mass spectrometry imaging enables spatial metabolomics of optimal cutting temperature compound (OCT)-embedded tumors

**DOI:** 10.1101/2025.11.05.686836

**Authors:** Joseph Monaghan, Nicholas Woytowich, Tian Zhao, Kiera Nguyen, Emily Mahony, Julian J. Lum, Kyle D. Duncan

## Abstract

Mass spectrometry imaging (MSI) is emerging as a powerful tool for uncovering the distribution of metabolites in the tumor microenvironment and studying tumor metabolism *in vivo*. However, to date, MSI of primary patient biobanked tissues contextualized by patient data has been limited to peptides, proteins, and glycans – with few examples for metabolites. This is because most biobanked fresh-frozen tissue required for spatial metabolomics is embedded in optimal cutting temperature compound (OCT), which introduces high-abundance polymeric interferents. Herein, we use nanospray desorption electrospray ionization (nano-DESI) to demonstrate the MSI of metabolites in OCT-embedded tissue. Metabolite coverage and sensitivity for prepared tissue mimetic homogenates embedded in OCT and an MSI-compatible material, carboxymethylcellulose (CMC), showed excellent agreement. We apply our ambient MSI workflow to detect changes in intratumoral methionine using a preclinical cancer mouse model undergoing adoptive T-cell therapy. Eight days after tumor incubation, lymphoma-bearing mice were maintained on a complete or methionine-restricted diet for 2 days. Nano-DESI MSI revealed a heterogeneous tumor microenvironment, with multiple methionine-cycle intermediates (S-adenosylmethionine, S-adenosylhomocysteine) and related metabolites, including known T-cell modulators (1-methylnicotinamide, polyamines) localizing to tumor subregions. Methionine-restricted tumors exhibited reduced methionine levels and elevated S-adenosylmethionine, relative to the control group. Overall, this work demonstrates spatial metabolomics on fresh-frozen OCT-embedded tissue, unlocking the wealth of information stored in primary tissue biobanks and consequently accelerating our understanding of cancer metabolism and treatment.

## 1. Introduction

Spatial metabolomics leverages mass spectrometry imaging (MSI) to visualize the abundance and localization of metabolites, providing information on an organism’s phenotypic response to biological or environmental stressors.^1–4^ MSI provides rich datasets reporting the spatial coordinates for a wide range of primary and secondary metabolites that are difficult/impossible to capture using conventional histology,^1,3,5–11^ and has been widely applied in fresh-frozen biospecimens, including analysis of drug/metabolite distributions,^12^ understanding developmental cascades,^13^ and as a probe to identify tumor/healthy tissue margin.^14–16^ Recent advances extend spatial metabolomics by combining the data with additional imaging modalities such as immunofluorescence or spatial transcriptomics.^17–20^ When co-registered, localized metabolism from MSI can be annotated by cell type or the local transcriptome, providing a multidimensional understanding of complex, heterogeneous tissues including the tumor microenvironment.^21–24^

A compelling application of MSI is in unraveling molecular mechanisms of cancer pathology, revealing insights into how the disease progresses or why treatments fail.^25^ Imaging retrospectively-collected tissue samples where data can be contextualized against known patient treatment history and outcome is particularly valuable.^26^ However, most biobanked tissues are embedded in paraffin wax (e.g. formalin-fixed paraffin-embedded [FFPE]) or optimal-cutting temperature compound (OCT).^10,27–29^ This challenges spatial metabolomics workflows, as embedding can delocalize metabolites or introduce interfering polymers that suppress ionization and contaminate the mass spectrometer.^30^ For MSI, fresh-frozen tissues with no embedding are preferred; however, this is not always possible for small and/or fragile tissues.^31^ Studies incorporating spatial metabolomics generally favor materials previously established to be compatible with MSI, such as carboxymethylcellulose (CMC) or gelatin, however, this requires MSI to be considered from the outset.^30^ Historically, tissues from mouse models and human patients have been archived in FFPE, OCT, or flash-frozen without matrix. Without a systematic understanding of the impacts of embedding matrices on small molecule metabolites, analyzing these embedded tissues with MSI has been considered impractical – locking away metabolic features that could inform valuable mechanistic insights on disease pathology.

Several groups have explored imaging biomolecules in FFPE tissues,^12,32,33^ however, the numerous rinse steps needed for tissue fixation can wash out and delocalize both polar and non-polar molecules. This limits the biological conclusions that can be obtained with MSI of small molecules in FFPE tissue.^7,27^ In contrast, OCT-embedding is performed on fresh-frozen tissues, providing a more promising alternative with less risk of metabolite delocalization. However, the MSI community has been resistant to imaging OCT-embedded tissues, as OCT contains ionizable polymers including polyvinyl alcohol and polyethylene glycol that contribute to chemical background and risks MS contamination.^12,27,28,34^ Despite these challenges, several groups have applied MSI to image biomolecules in OCT-embedded tissue.^23,28,35– 41^ Early work from Schwartz *et al*. compared rat liver tissues with and without OCT-embedding and noted fewer protein features in OCT-embedded tissue.^28^ Other groups have explored aqueous washing of OCT-embedded tissue to mitigate chemical background.^36,37,42^ While promising for some molecules, these studies focus on proteins or lipids and do not establish OCT compatibility for small molecule metabolites.

Herein, we use nanospray desorption electrospray ionization (nano-DESI) MSI to establish the viability of spatial metabolomics for OCT-embedded tissues. With contrived tissue homogenates, we show that comparable coverage and sensitivity can be achieved from both CMC- and OCT-embedded tissues. Imaging of an OCT-embedded brain revealed metabolite localization to expected anatomical features, suggesting the embedding process does not obfuscate spatial information. The developed nano-DESI MSI workflow was applied to a preclinical murine cancer model where tumor cells were implanted into immunocompetent mice and randomized into a control diet or methionine-restricted diet. For each experimental arm, matched tumors were bisected, with one half embedded in CMC and the other in OCT prior to imaging. The resulting metabolic perturbations caused by methionine restriction were compared between embedding media to evaluate if similar insights could be generated from OCT-embedded tissues. To our knowledge, this is the first study to comprehensively characterize the impacts of OCT-embedding on small molecule metabolites for MSI – demonstrating the capacity for wide coverage and to capture metabolite localization despite OCT interference.

## 2. Materials & Methods

### 2.1 Chemicals and standard solutions

HPLC-grade methanol, HPLC-grade water, and 88% formic acid were purchased from Fisher Scientific (Ottawa, ON). Rat brain (Sprague-Dawley) was purchased from Charles River Laboratories (Wilmington, MA, USA). Optimal Cutting Temperature (OCT) compound was purchased from Fisher Scientific (Cat #23-730-571). Carboxymethylcellulose (CMC) embedding material was purchased from Epredia (M-1 Embedding Matrix; Catalogue #: 1310; Kalamazoo, MI, USA). ^15^N-labelled amino acid internal standard mixture (Sigma-Aldrich; Cell free amino acid mixture 15N; 98 atom%, Oakville, ON) and lysoPC 19:0 (Avanti Research; Alabaster, AL) were included as internal standards at ∼3 µM for each amino acid and 5 µM for the lysoPC. The nanoDESI solvent was a 9:1 mixture of methanol:water, spiked fresh with 0.1% formic acid (*v/v*) on the day of imaging.

### 2.2 Preparation of rat brain homogenate

Rat brain homogenate was generated with ∼ 1 g rat brain added to a homogenizer microtube along with one 5-mm steel bead (Figure S1). Homogenization was carried out at 5.0 m/s for 50 seconds using a Bead Mill 4 Homogenizer (FisherBrand; Fisher Scientific, Ottawa, ON). After homogenization, material was withdrawn into the tip of a 1000 µL positive displacement pipette and subsequently wrapped in parafilm and frozen on dry ice for ∼30 mins. The end of the pipette tip was removed using a clean utility knife, and the remaining homogenate was ejected slowly onto cooled aluminum foil. The frozen homogenate was cut into ∼200µL sections for CMC embedding, OCT embedding, or LC-MS. To embed the homogenates, ‘rafts’ were constructed from aluminum foil and cooled on dry ice along with a small amount (2–3 drops) of embedding media (CMC or OCT). Homogenates were transferred onto the rafts via cooled forceps, and the respective embedding media were added until the homogenate was completely submerged. The embedded homogenates were left to freeze on dry ice for 60 minutes. Homogenate sections were then collected at 12 µm thickness via cryotome (Leica CM1850, Wetzlar, Germany) and plated directly to regular microscope slides (SuperFrost), before storage at –80 °C.

### 2.3 Murine tumor model

All animal studies were conducted in accordance with the Canadian Council on Animal Care guidelines and received approval from the University of Victoria’s Animal Care Committee (2023-019), University of British Columbia - BC Cancer Research Ethics Board (H25-00625) and University of Victoria Human Research Ethics Board (19-0067). Animal studies have been described in detail previously.^43^ Briefly, experiments were performed with the Animal Care Service (ACS) at the University of Victoria. Female B6 Thy1.1 mice (4 months old) were implanted with 10^6^ EG7-OVA tumor cells (Day 0). On Day 8, mice were infused with 6.5x10^5^ activated CD8^+^ OT-I T-cells through the tail vein and put on a control diet (0.86% methionine, Teklad Cat. #: TD230144) or a methionine-restricted diet (0.06% methionine, Teklad Cat. #: TD230293). Tumor volumes were measured twice a week, and mice were euthanized and tumors were harvested when tumor volumes exceeded 1500 mm^3^ or mice lost >20% of their initial body weight. Freshly isolated tumors were immediately embedded in CMC or OCT in aluminum foil ‘rafts’, flash-frozen in the vapor phase of liquid nitrogen using a CryoPod Carrier (Azenta Life Sciences) and stored at

−80 °C for approximately 18 months prior to imaging.

### 2.4 nano-DESI mass spectrometry imaging

nano-DESI imaging was performed on an Orbitrap Exploris 120 (ThermoScientific, San Jose, CA, USA) using a custom interface based on previous designs (Figure S2).^44,45^ Briefly, two fused silica capillaries (50 µm inner diameter, 150 µm outer diameter; Molex, Lisle, IL, USA) were positioned next to one another at an ∼90° angle. A 0.5 µL/min flow of spray solvent (9:1 methanol:water with 0.1% formic acid, 3 µM ^15^N-amino acid internal standards, and 5 µM LysoPC 19:0 internal standard) was delivered to the primary capillary, which was pneumatically aspirated through the secondary capillary using a controlled flow of UHP 5.0 N_2_ gas (1–3 L/min, SFC-6000D-5SLPM; Sensirion, Stäfa, Switzerland). For imaging, the capillary assembly was held fixed, and the tissue was rastered underneath using a precision XYZ stage (Zaber Technologies, Vancouver, BC, Canada). In contrast with previous nano-DESI imaging experiments, we only imaged tissue-associated regions by dropping the stage ∼500 µm below the capillaries for background or embedding material associated regions. To collect the images, the sample was scanned along the *x-*axis at 40 or 60 µm/sec, and stepped at 150 µm increments along the *y*-axis, resulting in pixel sizes of 20–30 µm (X) by 150 µm (Y), determined through the lateral movement of the sample and the MS duty cycle (X) and step size between scan lines (Y). The stages and mass flow controller were controlled using newly developed software written in Python 3.12, similar to previously described LabView software.^45^ The MS inlet was heated to 250 °C, and an ESI voltage of +3.4 kV (positive mode) was applied to the primary capillary. Full-scan data were collected at a target resolution of 120,000 at *m/z* 200. Images were acquired with a segmented full scan for the low and high *m/z* range, with windows of *m/z* 70-500 and 480-1000, respectively.

### 2.5 nano-DESI mass spectrometry imaging data processing

Vendor ‘.raw’ data files from MSI were converted to imzML using imzML Writer.^46^ All image files for this project are available as a METASPACE^47^ project at [Available upon Publication]. Subsequent processing of the imzML files was conducted using custom Python & R scripts to extract tissue-associated pixels based on an ROI mask, align *m/z* features across spectra/images, and visualize/pool the resulting data as ion images or volcano plots. Briefly, ion images were generated in imzML Scout with a 5ppm mass tolerance (unless otherwise specified). Average mass spectra were generated in Cardinal 3.6 (R Package).^48^ Volcano plots and log-log intensity plots were generated in Python 3.12 based on the tissue-associated, MS-aligned pixel spectra or average spectra generated using Cardinal. PCA was carried out using MetaboAnalyst 6.0 using the ‘Statistical Analysis [metadata table]’ module.^49^ Average spectra for the lymphoma tumor images were input with associated metadata, and spectra were sum-normalized (per-sample), then mean-centred and pareto-scaled (across-samples) prior to PCA. To focus the PCA on endogenous metabolic features and mitigate interference from the OCT chemical background, only *m/z* features observed in both the CMC- and OCT-embedded tumors were included in the analysis.

### 2.6 Metabolite Annotation

Where possible, metabolites were annotated based on LC-MS analysis of the same tissue sample used for imaging. MS and LC models, column and gradient details, and instrument parameters are summarized in Tables S1 and S2. Metabolites labelled ‘putative’ could not be conclusively identified due to an imperfect MS/MS match or multiple peaks in LC trace. Those without this label exhibited excellent MS/MS matching to spectral databases (HMDB, MassBank)^50,51^ or relevant literature^52,53^ and one peak in the LC-MS chromatogram (Figures S3 and S4). If metabolites were not detected by LC-MS, annotations were confirmed using on-tissue MS/MS (Figure S5). To prepare samples for LC-MS, rat brain homogenate or OCT-embedded tumor was weighed into a 4 mL GC vial. Methanol was added to the vial until the tissue concentration was ∼35 mg tissue/mL MeOH and sonicated for 60 mins at room temperature (∼20 °C), before being left to settle on the bench for a further 90 mins. The supernatant was collected and transferred to an Eppendorf tube and spun at 2g for 6 mins. The supernatant was collected again and transferred to a 2 mL amber GC vial and stored at –20 °C prior to LC-MS analysis.

## 3. Results & Discussion

### 3.1 Comparable metabolomic coverage & sensitivity is achieved from OCT-embedded tissues

To measure the impact of OCT-embedding on nano-DESI sensitivity and coverage, we prepared a rat brain homogenate embedded in both OCT and CMC (Figure 1A and 1B). The brain was homogenized, loaded into the tip of a positive displacement pipette, and frozen on dry ice (Figure S1). Once frozen, the end of the pipette tip was cut and the homogenate was split for CMC or OCT embedding. This approach ensured that the distribution and abundance of metabolites are the same for both embedding materials, which isolates the effect of the embedding material. The average mass spectra for both homogenates are shown in Figure 1C, with OCT-associated peaks highlighted in red (OCT spectrum available in Figure S6). Panels 1D and 1E compare median *m/z* intensities for low- and high-mass range features observed in the CMC-embedded homogenate. Grey data points are consistent across embedding materials (< 2-fold change in signal), blue points are higher in the CMC, and red points are higher in the OCT. Most features below *m/z* 480 exhibit very similar intensities in both homogenates, demonstrating that small molecule metabolites can be easily detected in OCT-embedded tissue.

**Figure 1.**
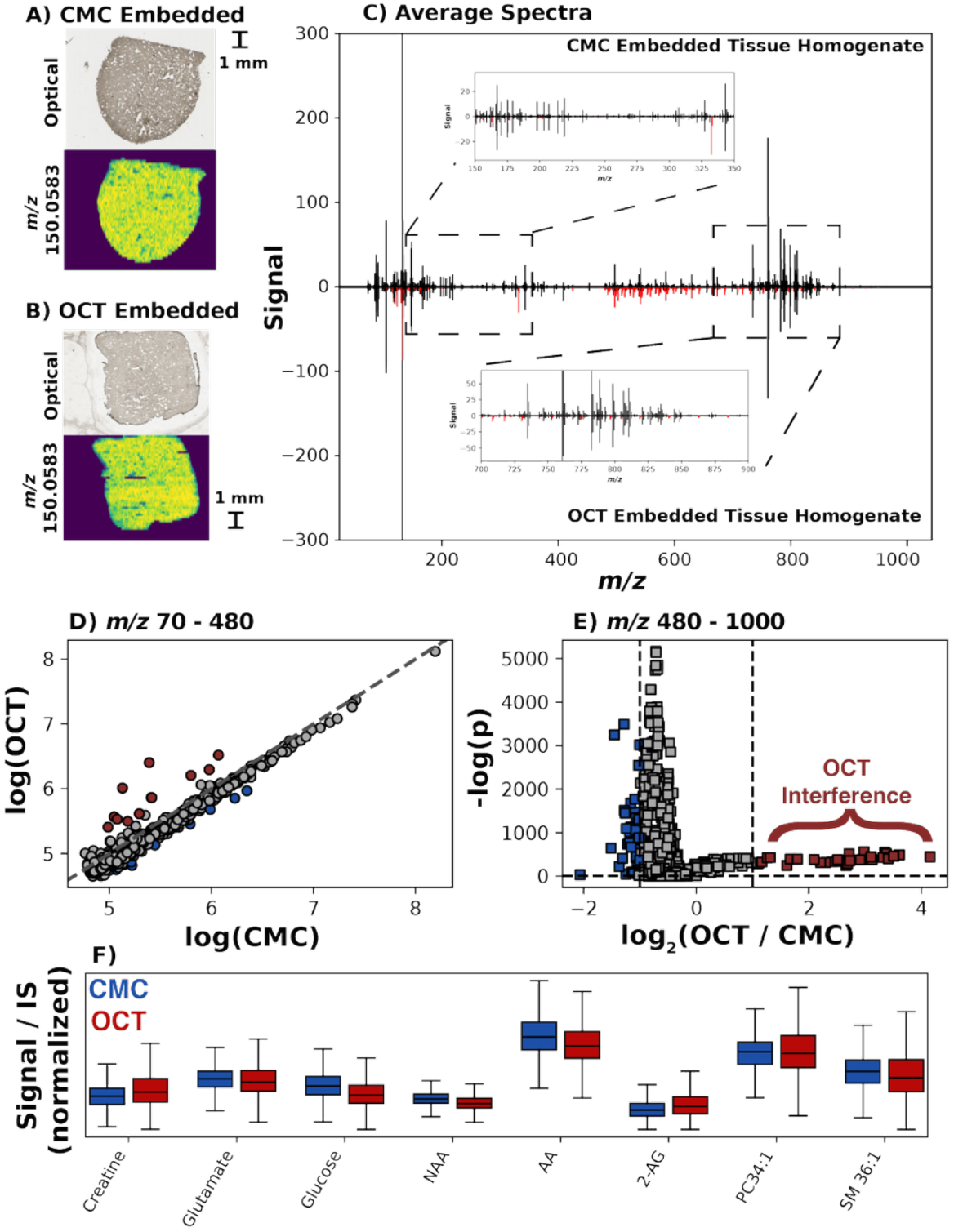
nano-DESI MSI comparison of brain homogenate with CMC or OCT-embedding. A & B) Optical and ion image for the CMC-embedded (A) and OCT-embedded (B) tissue homogenate. C) Average mass spectra in positive ion mode for each homogenate. Peaks in red indicate *m/z* associated with OCT [M+H], [M+Na], or [M+K] ions. Insets show optical image and TIC-normalized ion images for methionine (*m/z* 150.0583) for each embedding material. D) log-log intensity plot for median peak intensities in the CMC vs. OCT embedded tissues. The dotted line indicates a 1:1 relationship. Points in red indicate *m/z* that are measured at greater than 2-fold higher abundance in the OCT-embedded homogenate. E) Volcano plot for *m/z* median intensities in the OCT vs. CMC data. Points in red and blue indicate *m/z* that are greater than 2-fold abundance in either the OCT or CMC embedded homogenate, respectively. F) Boxplots comparing signal intensities for the OCT and CMC homogenate after normalization to a ^15^N-methionine (low mass) or LysoPC 19:0 (high mass) internal standard included in the nano-DESI solvent. To allow visualization of all metabolites on the same scale, internal standard normalized intensities are normalized to the maximum observed value. NAA: N-acetyl-aspartic acid [putative ID]; AA: Arachidonic acid; 2-AG: 2-arachidonoylglycerol [putative ID]. PC34:1 and SM 36:1 represent a mixture of isomeric lipids.

However, significant chemical background was observed in the *m/z* 400–700 range (peaks in red; Figure 1C). Some of these OCT peaks are in close *m/z* proximity with endogenous metabolites and lipids, as several ‘endogenous’ features (*i.e*. observed in CMC homogenate) exhibited higher intensity in the OCT homogenate (*n* = 31; Figure 1E). Of these, 28 of 31 are within 5ppm of a *m/z* observed in the pure OCT spectrum or an analogous [M+Na] or [M+K] adduct, suggesting interference from the chemical background. Other features exhibit significantly lower intensity in the OCT-embedded tissues (Figure 1E; *n* = 40 reduced features, Table S3). This reduction is particularly prevalent between *m/z* 480–650 (25 of 40), which may be driven by space-charge effects or artifacts from the vendor eFT pre-processing, given their proximity to intense OCT background (Figure S6).^54^ While lower absolute abundance is observed with OCT at this mass range, the effect can be compensated for by normalizing to an internal standard in the nano-DESI solvent (Figure S7). This is apparent when comparing targeted metabolites, where highly similar distributions (Figure 1F) are observed for small molecules (creatine, glutamate) and larger lipid species (PC34:1 [putative], SM 36:1 [putative]). In general, caution should be exercised when interpreting non-targeted results from OCT-embedded tissues, and confident annotation based on MS/MS or LC-MS is recommended to rule out interference from OCT, particularly given that OCT peaks may shift in relative adduct abundance or charge state over an image. Overall, these homogenate experiments demonstrate that nano-DESI MSI achieves similar metabolic coverage and sensitivity in OCT-embedded tissues, particularly for small molecule metabolites below ∼ *m/z* 450.

### 3.2 Metabolites and lipids are conserved to substructures in OCT-embedded Tissues

While tissue homogenates showcased the feasibility of measuring metabolites from OCT-embedded tissues, they contain no defined substructures. To evaluate if metabolite localization(s) are preserved, we embedded an intact rat brain in OCT and imaged sagittal sections with nano-DESI (Figure 2). Encouragingly, the OCT background is confined to the outer layer of the tissue. No appreciable delocalization was observed, with several lipids exhibiting well-defined spatial patterns based on major anatomical features of the brain. For example, PC O-36:2 is localized to the corpus callosum and cerebellum. Similar patterns were observed for other metabolites, including PC O-38:6, 2-arachidonylglycerol (putative ID), and arachidonic acid.

**Figure 2.**
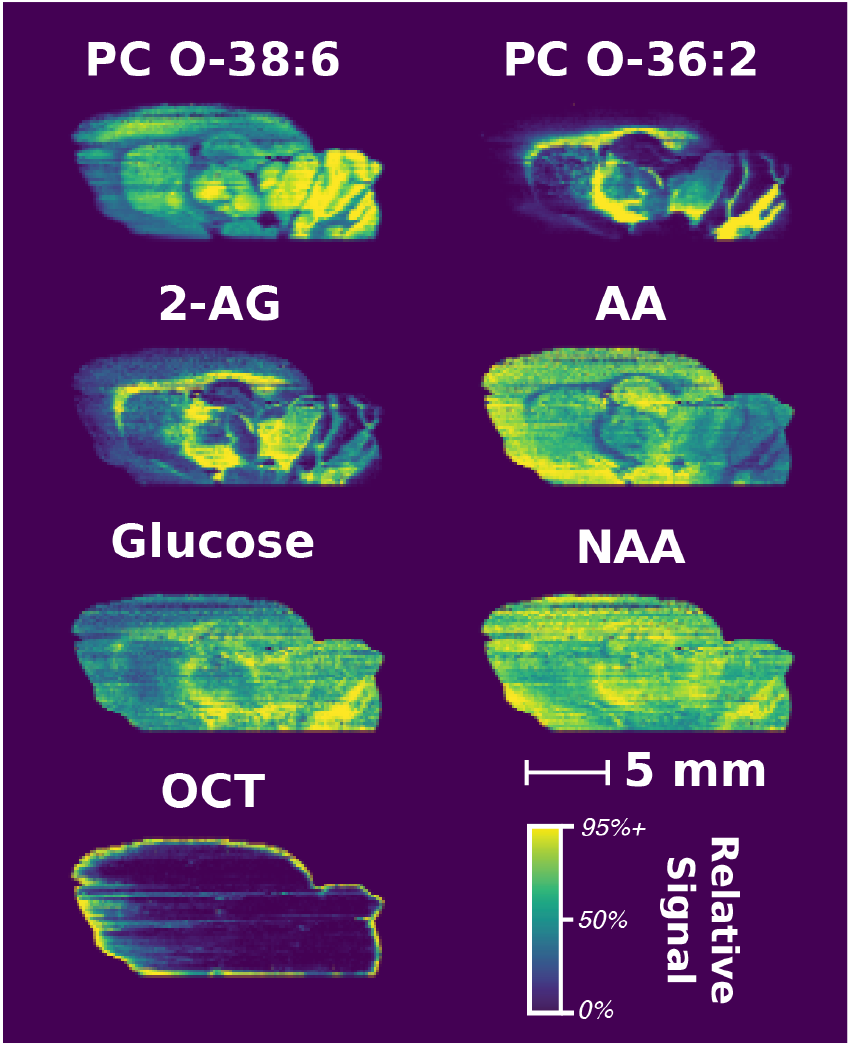
Ion images observed for an OCT-embedded brain cut along the sagittal plane. Ion images are TIC-normalized and generated with a 5ppm tolerance with hot pixels removed by setting the top of the color scale to the 95^th^ percentile. PC O-38:6 was monitored at *m/z* 806.5702, PC O-36:2 at *m/z* 772.6215, 2-AG (2-arachidonylglycerol; putative ID) at *m/z* 417.2402, AA (arachidonic acid) at *m/z* 327.2300, glucose at *m/z* 181.0707, NAA (N-acetylaspartic acid; putative ID) at *m/z* 176.0559, and OCT at *m/z* 752.3876.

Based on several studies reporting reduced OCT background after an aqueous washing step, a serial brain section was imaged after gently rinsing in DI water (60s + 30s in fresh DI). This step reduced OCT residue in the optical image of the tissue (Figure S8), and PC O-38:6 and PC O-36:2 showed similar localization with and without washing (Figures 2 and S9). However, small molecules were substantially reduced in the washed tissue (Figure S10). Further, these metabolites were redistributed to different regions of the brain. For example, glutamate is localized to the cortex in the unwashed tissue but appears predominantly in the cerebellum region after washing. Gamma-aminobutyric acid (GABA) shows a similar redistribution, with high abundance in two localized regions in the unwashed tissue, but is only observable in the cerebellum after washing. OCT features were also redistributed during the washing step, with *m/z* 752.3876 dispersed throughout the entire post-wash tissue (Figure 2; Figure S9). Together, these results suggest that while an aqueous washing step may improve performance for hydrophobic lipid species or proteins, it is incompatible with imaging of water-soluble metabolites.

Based on the OCT observed at the borders of the brain tissue, it was necessary to characterize the exchange of OCT and metabolites at the embedding material/tissue interface. Full line scans (*i.e*., embedding material and tissue) were collected for model lymphomas embedded in OCT or CMC.^43^ The optical image was manually aligned with MS data based on the signal for PC 34:1 (Figure 3A-D).^55^ As expected, no OCT is observed in the CMC-embedded tumor (3H), and OCT penetrated several mm into the tissue for the OCT-embedded tumor (3G). For PC 34:1, the [M+H], [M+Na], and [M+K] adducts appear at the tissue border for both CMC and OCT. However, glutamate [M+H] is observed 200–500 µm before the start of the tissue for both embedding matrices, suggesting diffusion beyond the tissue boundary during the embedding and/or sectioning process. Interestingly, glutamate [M+Na] and [M+K] adducts are only observed within the tissue border for OCT, which may be due to abundant OCT polymers outcompeting for K^+^ and Na^+^ ions. Diffusion of hydrophilic metabolites is consistent with previous observations by Li et al., where hydrophilic metabolites diffused further from the tissue border compared to hydrophobic lipids in embedded tissues.^30^ These data indicate care should be given when interpreting results near the embedded tissue border, as diffusion can drive metabolite delocalization.

**Figure 3.**
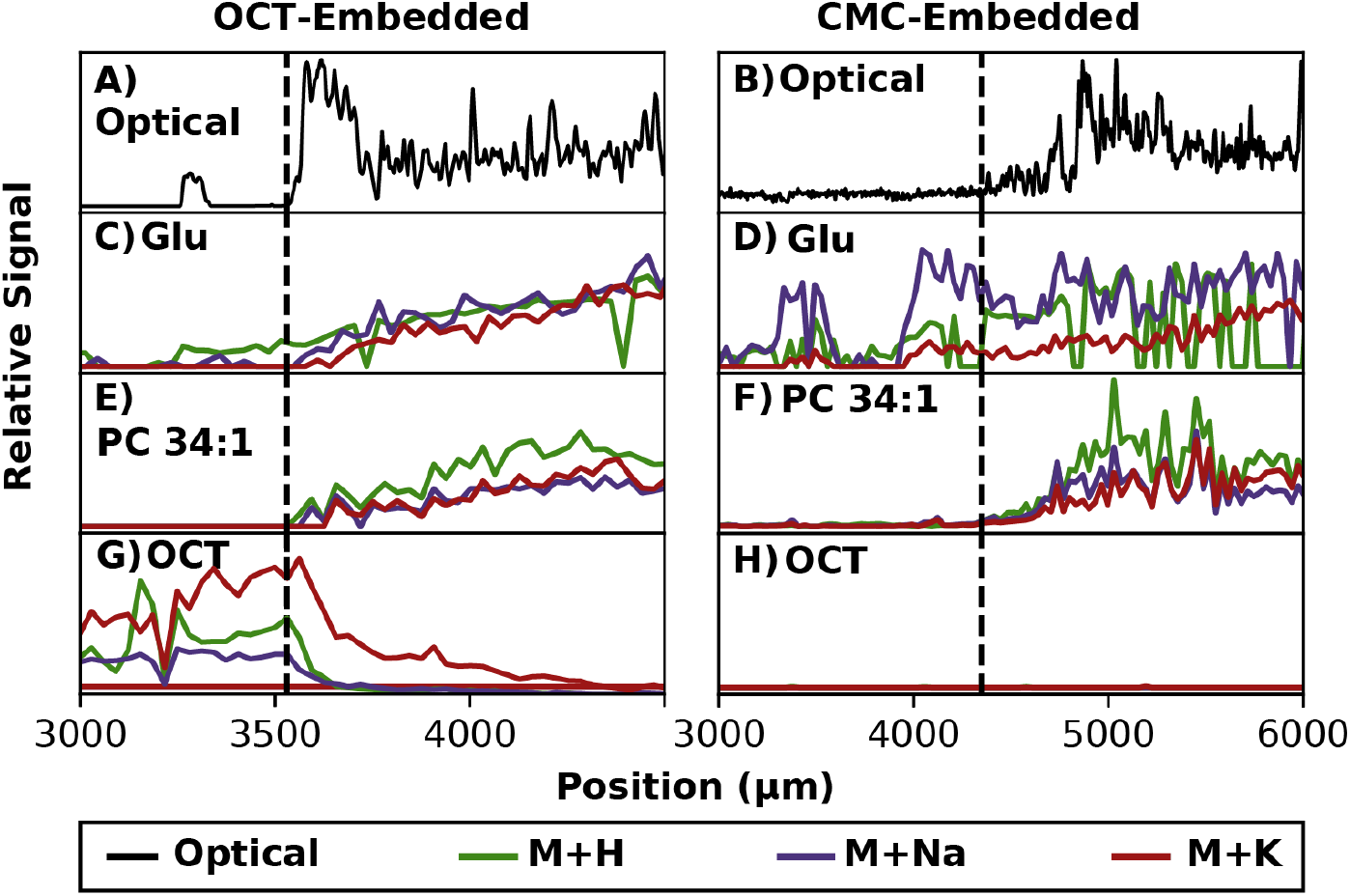
nano-DESI line scans demonstrating exchange of OCT embedding material and metabolites between tissue and the embedding medium interface. Glu: Glutamate [M+H: *m/z* 148.0610], PC 34:1 [M+H: *m/*z 760.5851], OCT [*m/z* 505.9745]. The optical trace was extracted from a slide scanner image of the whole tissue, averaging the grayscale intensity across a ∼150 µm transect of the tissue corresponding to where the line scan was performed.

### 3.3 Comparable metabolic perturbations are observed for tumors embedded in OCT and CMC

To demonstrate a proof-of-principle application of nano-DESI MSI for OCT-embedded tissues, we investigated methionine cycle intermediates and related metabolites in OCT-embedded lymphoma tumors. The methionine cycle is emerging as a key regulator of T-cell function in the tumor microenvironment.^43,56,57^ Moreover, these metabolites span a range of physiological concentrations and molecular properties (Table S4),^51^ with some very abundant (methionine; 20–30 µM in blood) and others 10–1000 times lower (SAH: 0.01–0.5 µM in blood; MNA: 0.007–0.85 µM in blood). Spatially resolving the distribution of these metabolites could help design cellular-based therapies with improved metabolic fidelity and *in vivo* efficacy.^7,57^ Figure 4 shows the methionine cycle with ion images for each intermediate. Methionine and nicotinamide are homogeneous throughout the tissue, whereas SAM, SAH, and MNA localize to subregions of the tumor. Homocysteine was not detected, likely due to a poor ionization efficiency in positive ion mode, which could be overcome using reactive nano-DESI strategies such as functionalizing the thiol with a charged pyridinium tag.^58^ The spatial trends for the methionine cycle intermediates were observed across multiple technical replicates (*n* = 3; Figure S11) of the same tumor as well as biological replicates (*n* = 3; Figure S12). Overall, these results indicate that nano-DESI MSI is an effective tool for imaging small molecule metabolites, even in OCT-embedded tumors.

**Figure 4.**
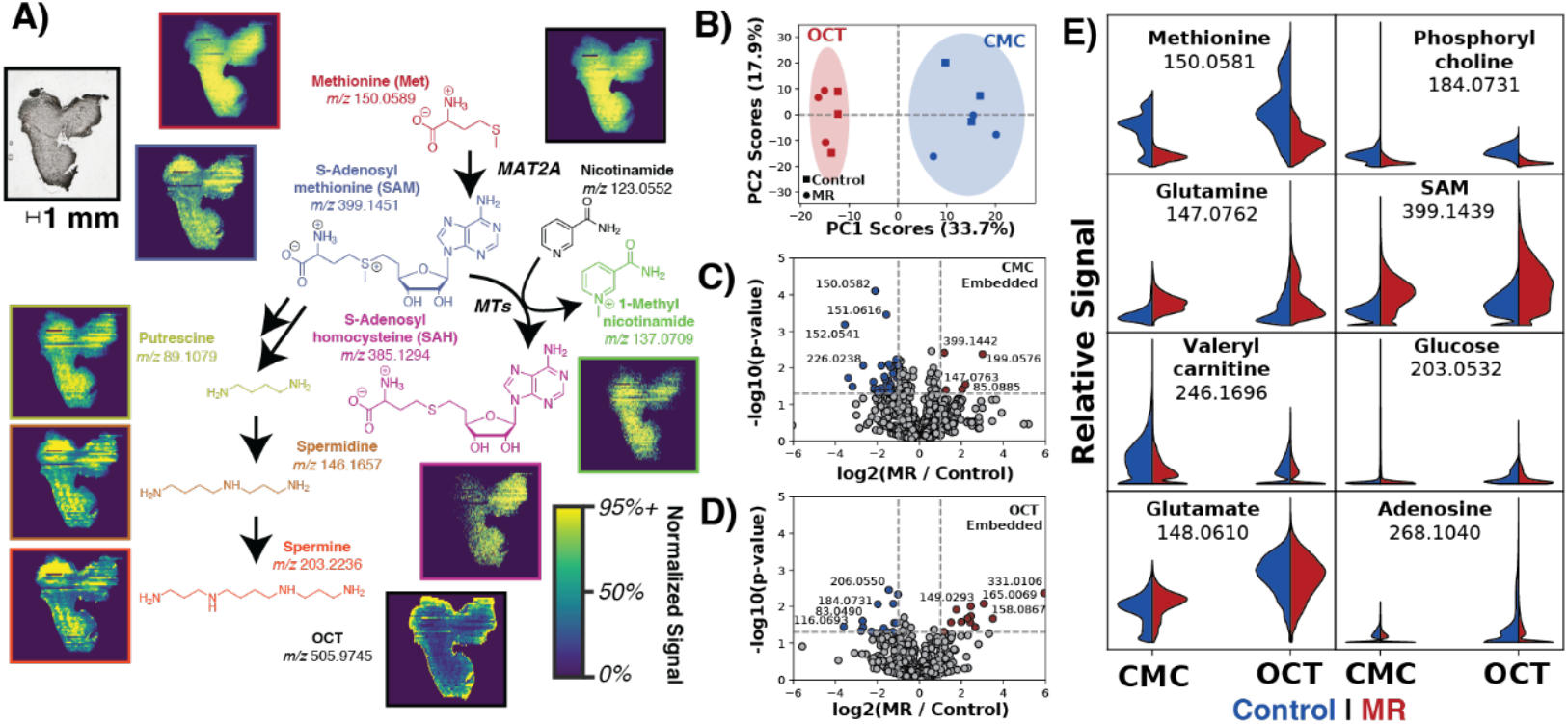
nano-DESI MSI of OCT-embedded tumors. A) Ion images for methionine cycle intermediates and related metabolites in an OCT-embedded tumor section. Hot pixels are removed by setting the color scale cutoff at the 95^th^ percentile of each image’s dataset. Images are generated with a 5ppm tolerance to the target *m/z*. B) PCA scores plot comparing average full scan data for matched OCT and CMC embedded tumor sections collected from lymphoma-bearing mice that were maintained on either a complete (control) or methionine-restricted diet (mean-centred & pareto-scaled). C) and D) Volcano plots comparing feature abundance between control and MR conditions for CMC-embedded and OCT-embedded tissues, respectively. E) Violin plots comparing whole-tissue pixel intensity distributions for select features, demonstrating similar relative distributions for both CMC and OCT.

A fundamental question for MSI of OCT-embedded tissues is if biological insights can be generated, especially compared with embedding materials considered MSI-compatible such as CMC. To assess this, mice bearing lymphoma tumors were adoptively transferred with tumor antigen-specific T cells and immediately fed *ad libitum* a methionine-restricted diet (*n* = 3) or control diet (*n* = 3) for two days. These six tumors were subsequently collected, split in half, and embedded in either OCT or CMC (12 images total). As expected, tumors grown under methionine-restricted conditions exhibited lower methionine abundance than control tumors across both embedding materials (Figures S13 and Zhao *et al*.).^43^ Using Cardinal^48^ and MetaboAnalyst,^49^ we generated a PCA plot from the aligned, average mass spectra of each tumor, revealing clear separation of OCT- and CMC-embedded tumors (Figure 4B). The loadings plot indicated low *m/z* features (104.1069, 132.1018, 76.0394) are most associated with OCT embedding, while higher *m/z* features (204.1228, 465.2613) are most associated with CMC (Figure S14). Volcano plots comparing methionine-restricted to control tumors revealed 29 and 28 features significantly differing at the *p* < 0.05 and absolute log_2_(fold-change) > 1 level for the CMC- and OCT-embedded tissues, respectively (Figure 4C/4D). These images are not perfectly paired, as they are collected from different faces of the same tumor, representing areas with potentially differing metabolite abundance. The data reflect this, as the identities of statistically significant features differ between CMC/OCT. However, the overall trends are consistent (Figure S15; Table S5). This is particularly apparent when the pooled, pixel-wise distribution of signal intensities is compared for specific features across the dataset (Figure 4E). For example, methionine, valerylcarnitine, and phosphorylcholine all appear depleted in the methionine-restricted tumors. Other metabolites appear slightly increased with methionine restriction, such as glutamine and SAM, while glucose, glutamate, and adenosine are largely unchanged with methionine restriction. Overall, the distribution of these metabolites is remarkably similar between the two embedding materials, suggesting comparable biological conclusions can be drawn from both CMC and OCT-embedded tissues.

While the results herein demonstrate the capability of MSI to localize metabolites (Figure 2) and extract differences in metabolite levels (Figure 4) in CMC and OCT-embedded tissue, fresh-frozen, unembedded tissues remain the gold standard. Factors such as tissue fragility and size (*i.e*., small tissues) often necessitate the use of embedding materials to maintain tissue morphology during storage and sectioning. Regardless of the MSI modality, metabolite exchange at the embedding material**/**tissue interface can confound interpretation. Further, while biobanks strive to collect and store high-quality tissues, in clinical environments, patient care is the priority and differences in tissue handling before or during embedding could shift the extent of exchange with embedding material on a tissue-to-tissue basis. Despite these limitations, the ability to image small molecule metabolites in OCT-embedded clinical samples demonstrated here is of enormous value to test hypotheses and develop an understanding of disease pathology with retrospectively collected tissue specimens.

## 4. Conclusions

This study demonstrates that nano-DESI can be used to image metabolites in OCT-embedded tissue samples. Tissue homogenates embedded in an MSI-compatible material, CMC, were compared to OCT-embedded tissue, which revealed excellent agreement for small molecule metabolites below *m/z* 450. For larger metabolites (> *m/z* 450), chemical background from the OCT is more prevalent and suppresses endogenous peak intensities. However, many lipids are still observed at reasonable signal-to-noise ratios, and internal standards can correct for some variance. OCT-embedding retained metabolite localization to defined tissue substructures that matched anatomical regions for both lipids and small molecule metabolites, where previously published washing steps led to substantially reduced signal intensities and delocalization of metabolites. Our workflow was applied to measure methionine cycle metabolites in both OCT- and CMC-embedded tumor sections collected from tumor-bearing mice fed a complete or methionine-restricted diet. While PCA unsurprisingly identified qualitative differences between OCT and CMC sample sets, similar relative abundances and statistically significant features were observed across both embedding materials. Overall, this work demonstrates that OCT-embedded tissues can be imaged with nano-DESI MSI to uncover the distribution of small molecule metabolites. Future application of our workflow to retrospectively collected biobanked tissue will enable MSI-based spatial metabolomics data to be contextualized with known patient outcomes/phenotypes and clinical data.

## Acknowledgments

We gratefully acknowledge the University of Victoria and Vancouver Island University for ongoing support for infrastructure and student researchers. This work was funded through the Natural Sciences and Engineering Research Council of Canada (NSERC; RGPIN-2022-03696), the Terry Fox Research Institute New Frontiers Program Project and the Lotte and John Hecht Memorial Foundation (Program Project Grant #1125), and MITACS Accelerate (IT39215). Infrastructure for the nano-DESI apparatus was funded through the Canadian Foundation for Innovation John R. Evans Leaders Fund (CFI JELF 43810). Salary support for J.M. and K.D.D. was provided through a trainee and scholar award (respectively) from Michael Smith Health Research BC. We acknowledge the Centre for Health and Environmental Mass Spectrometry (CHEMS) for access to MS instrumentation.

## Conflict of Interest

All authors have no conflicts of interest to disclose.

## Ethics Statement

Animal studies were carried out in accordance with the Canadian Council on Animal Care guidelines with approval from the University of Victoria’s Animal Care Committee (2023-019), University of British Columbia - BC Cancer Research Ethics Board (H25-00625) and University of Victoria Human Research Ethics Board (19-0067).

## Data Availability Statement

Imaging data is publicly available on METASPACE at [Available upon Publication].

## Notes

### Competing Interest Statement

The authors have declared no competing interest.

