## Supplemental Information for "Ambient mass spectrometry imaging enables spatial metabolomics of optimal cutting temperature compound (OCT)-embedded tumors"

**Table of Contents:**


*Figure S1.* 3

*Figure S2.* 4

*Figure S3.* 5

*Figure S4.* 6

*Figure S5.* 6

*Figure S7.* 7

*Figure S8.* 8

*Figure S9.* 8

*Figure S10.* 9

*Figure S11.* 10

*Figure S12.* 11

*Figure S13.* 12

*Figure S14.* 13

*Figure S15.* 13

*Table S1.* 14

*Table S2.* 14

*Table S3.* 15

*Table S4.* 16

*Table S5.* 17

*References:* 18

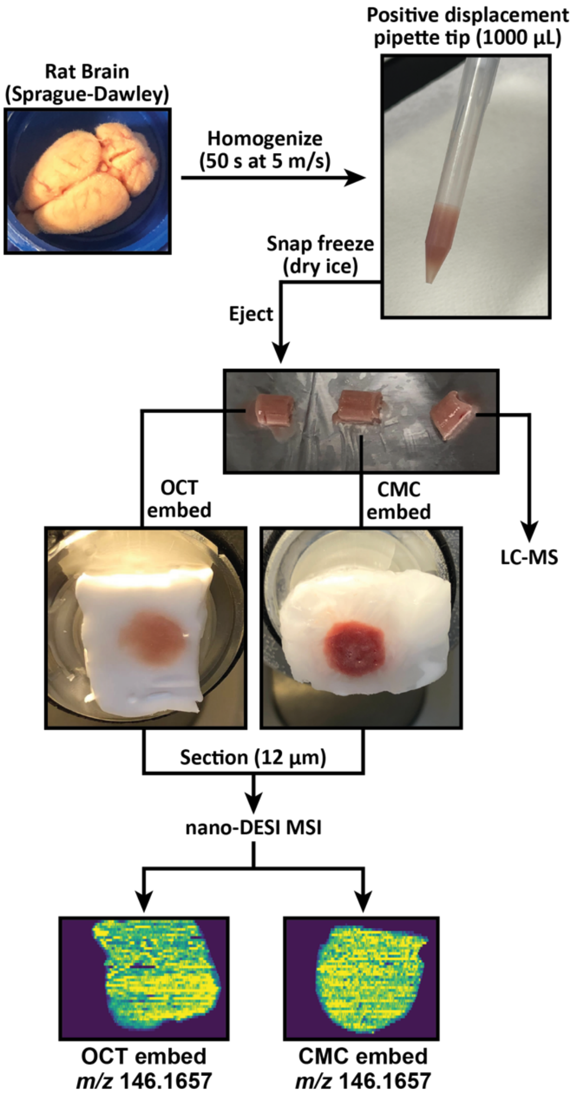


**Figure S1.** Overview of OCT Tissue preparation for tissue homogenate.

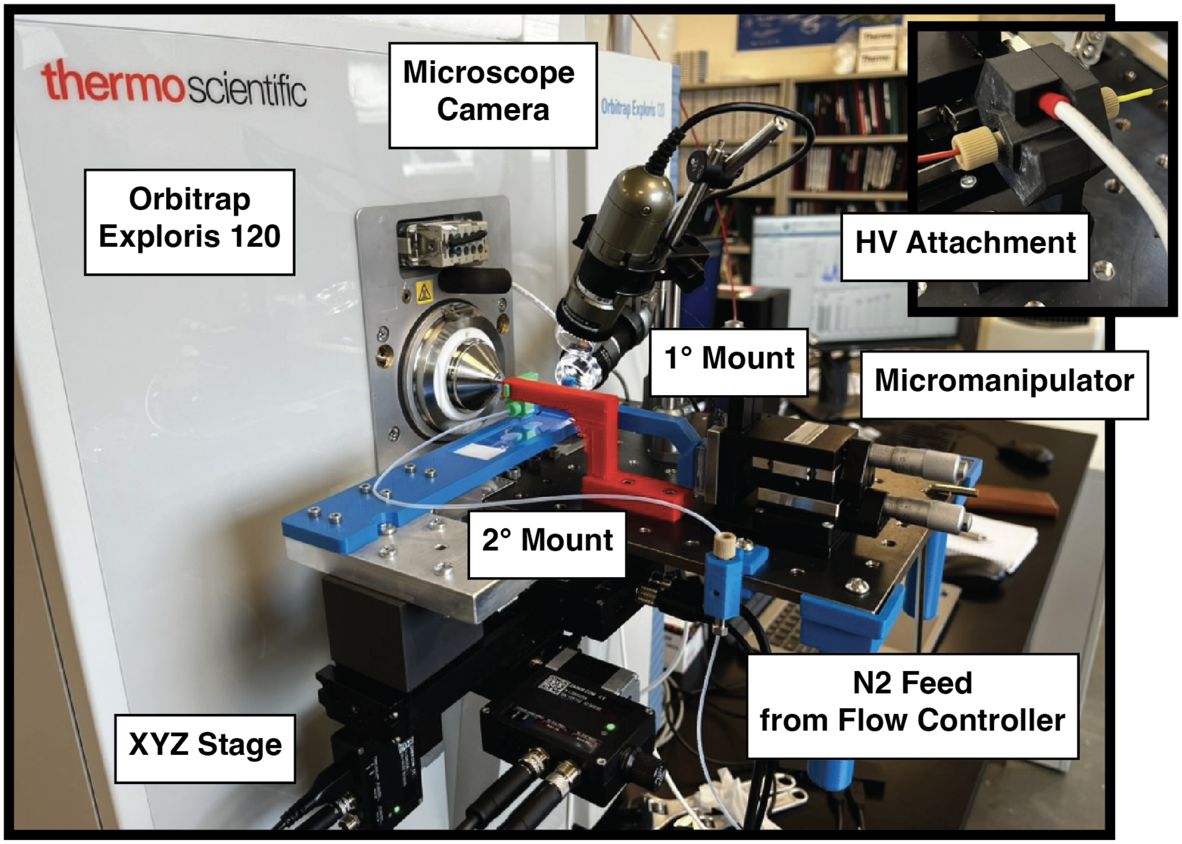


**Figure S2.** Photographs of the nano-DESI apparatus and 3D-printed capillary mounts.

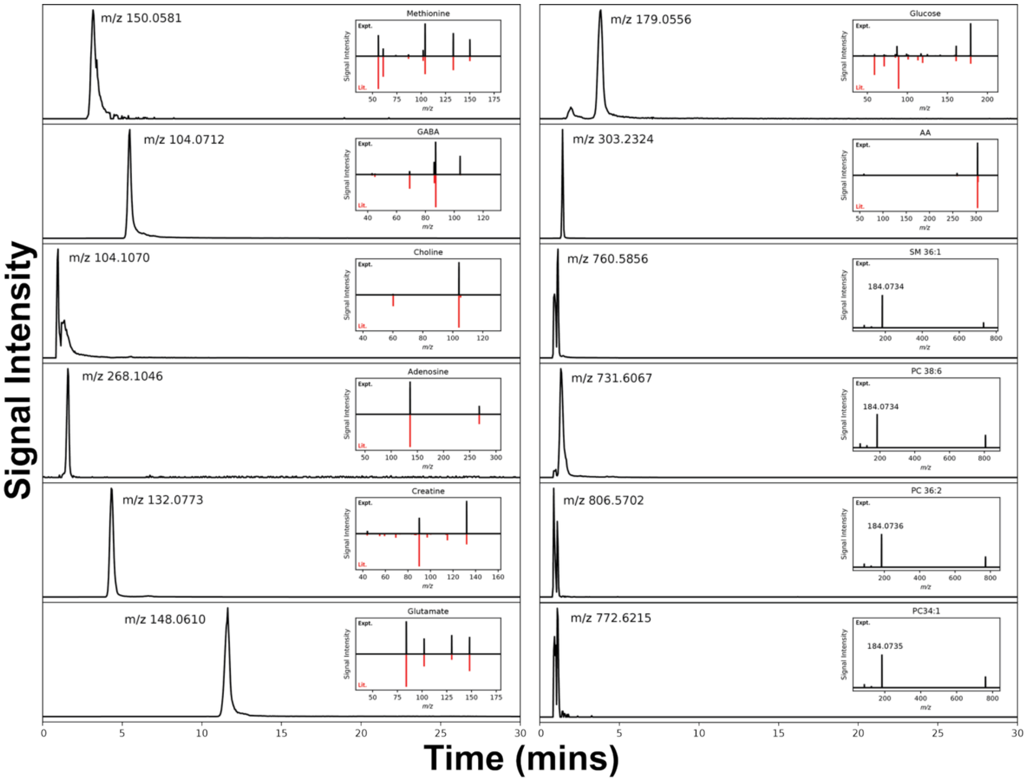


**Figure S3.** LC-MS/MS results for a methanolic rat brain extract. Chromatograms are exported with a 10ppm tolerance window. Literature spectra were retrieved from HMDB,[1] Massbank,[2] and relevant primary literature.[3] Experimental parameters for the LC run are available in Tables S1 and S2.

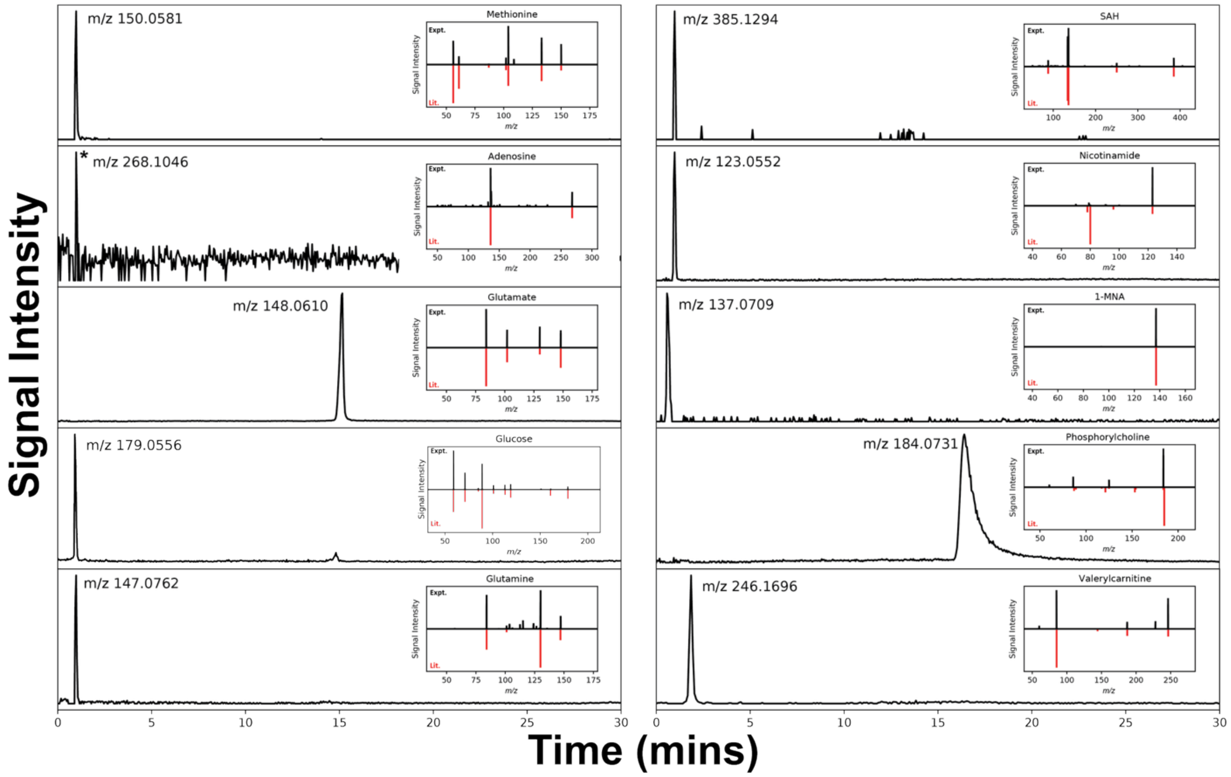


**Figure S4.** LC-MS/MS results for a methanolic extract of a control tumor sample. Chromatograms are exported with a 10ppm tolerance window. Literature spectra were retrieved from HMDB,[1] Massbank,[2] and relevant primary literature.[3,4] Experimental parameters for the LC run are available in Tables S1 and S2.

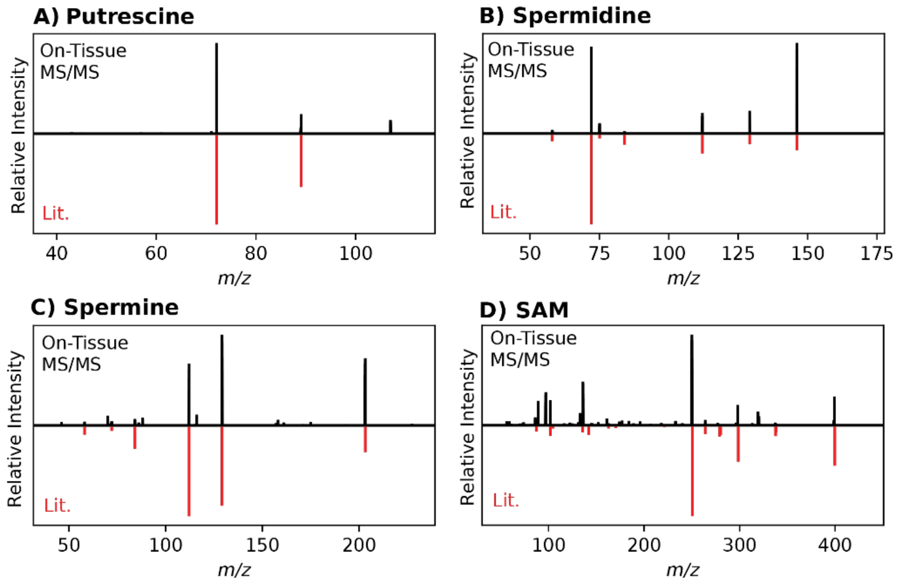


**Figure S5.** On-tissue MS/MS results compared with literature product ion scans for a control tumor sample. Literature product ion scans were retrieved from Massbank.[2]

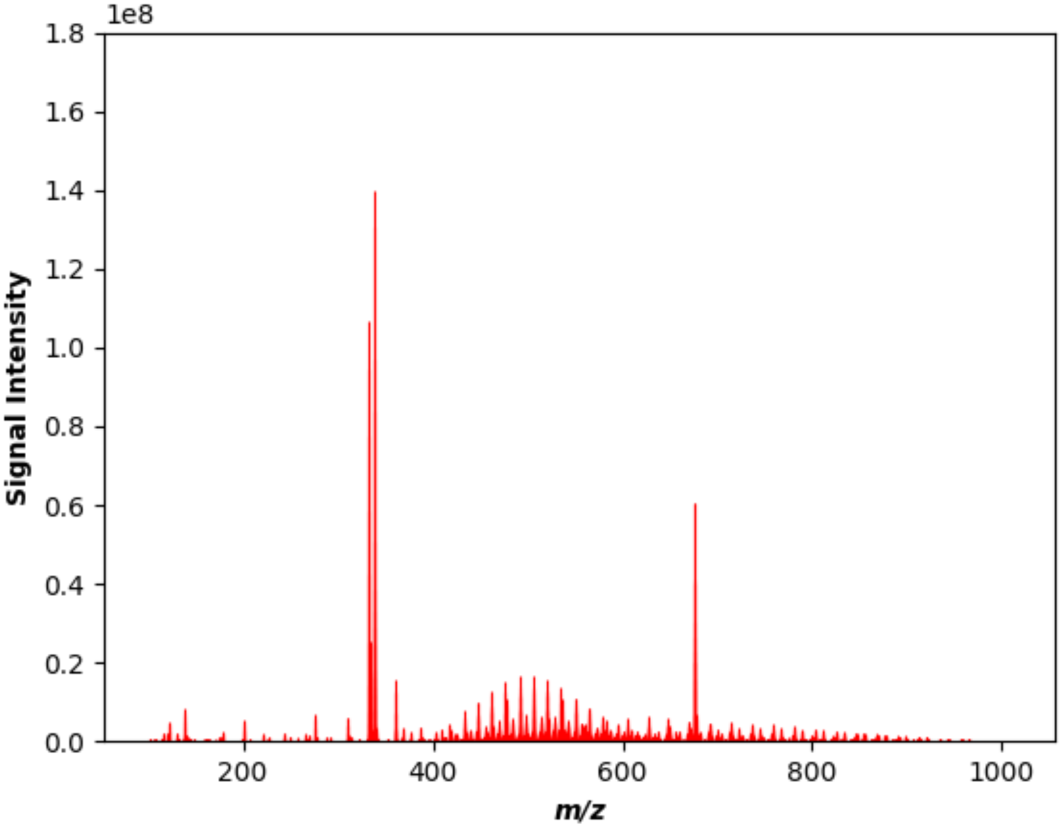


**Figure S6.** Average OCT spectrum collected from direct infusion of a 150ppm solution of OCT in 50:50 MeOH:Water.

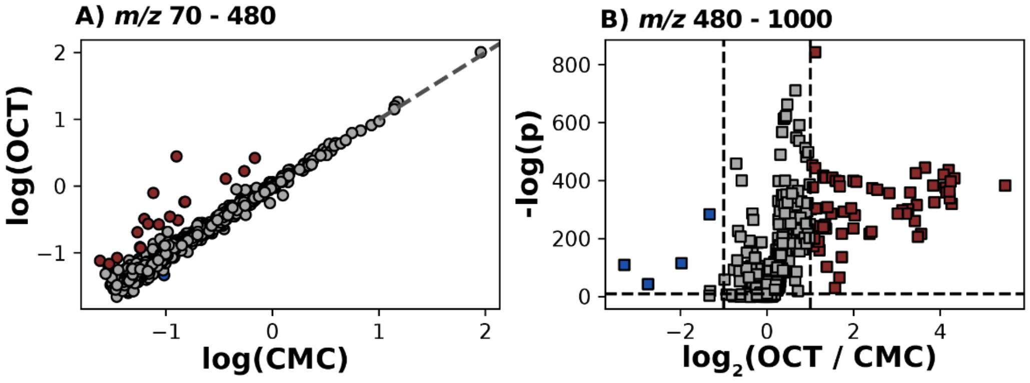


**Figure S7.** Nano-DESI MSI comparison of OCT- and CMC-embedded homogenates with internal standard normalization. $^{15}$N-methionine and lysoPC 19:0 were used as internal standards for the *m/z* 70-480 and 480-1000 ranges, respectively. Red markers indicate features that are > 2-fold more abundant in the OCT, while blue markers indicate peaks that are > 2-fold more abundant in the CMC.

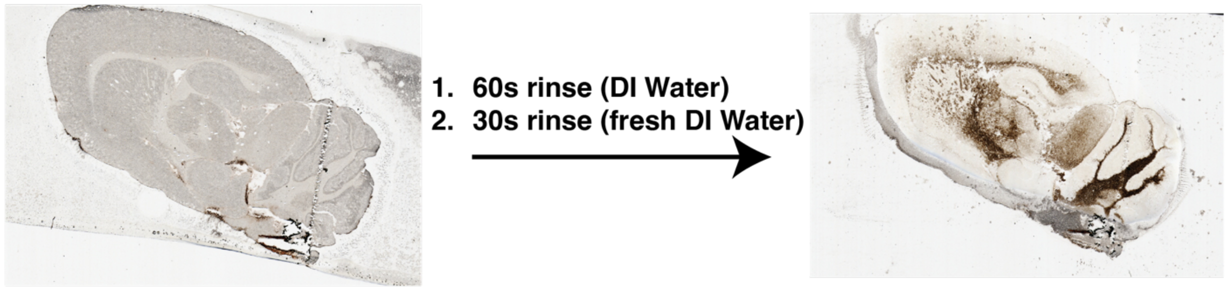


**Figure S8.** Before- and after-washing optical images for the sagittal brain section.

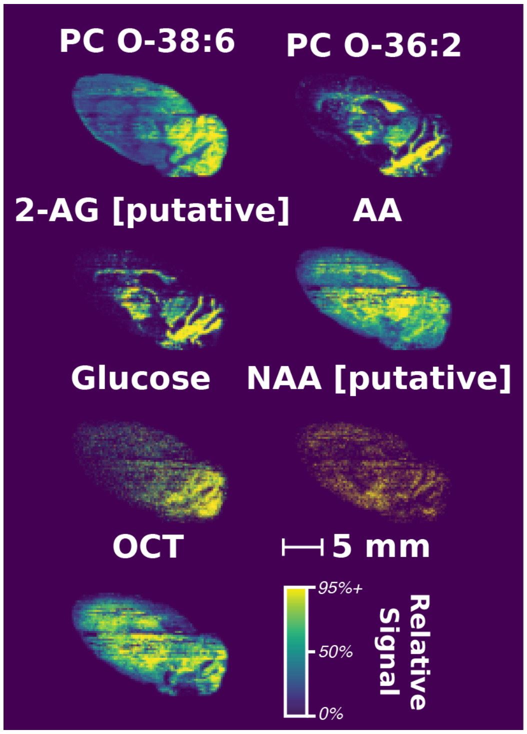


**Figure S9.** Delocalization and increased OCT background in ‘washed’ section of OCT-embedded rat brain (Sagittal). Ion images are TIC-normalized and generated with a 5ppm tolerance. Hot pixels were removed by setting the top of the color scale to the 95$^{th}$ percentile. PC O-38:6 was monitored at *m/z* 806.5702, PC O-36:2 at *m/z* 772.6215, 2-AG (2-arachidonylglycerol; putative ID) at *m/z* 417.2402, AA (arachidonic acid) at *m/z* 327.2300, glucose at *m/z* 181.0707, NAA (N-acetylaspartic acid; putative ID) at *m/z* 176.0559, and OCT at *m/z* 752.3876.

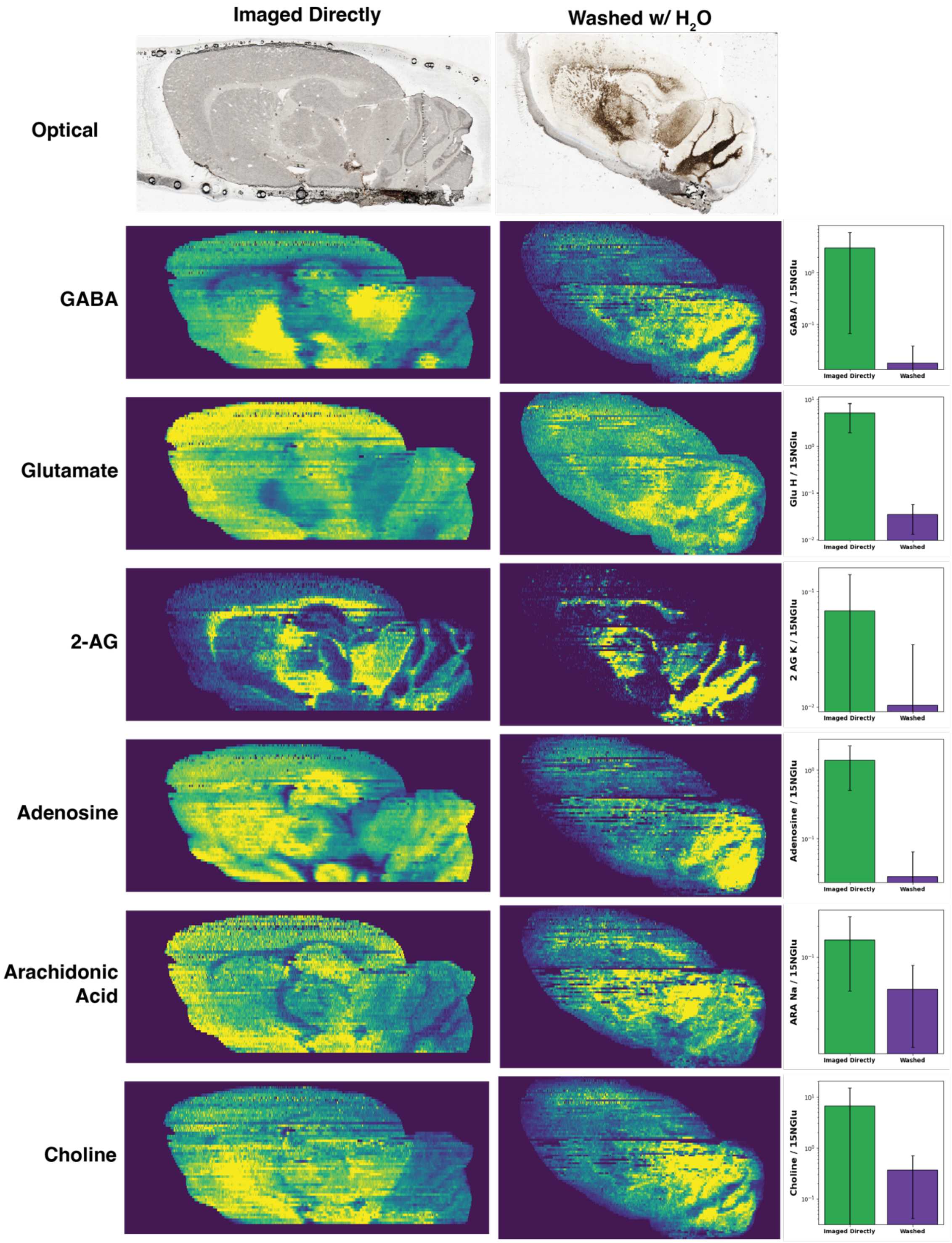


**Figure S10.** Comparison of Sagittal Rat brain section for a series of metabolites imaged directly (left) or after a washing step (right). In general, lower signal intensities are observed after washing, alongside delocalization and/or redistribution of several metabolites.

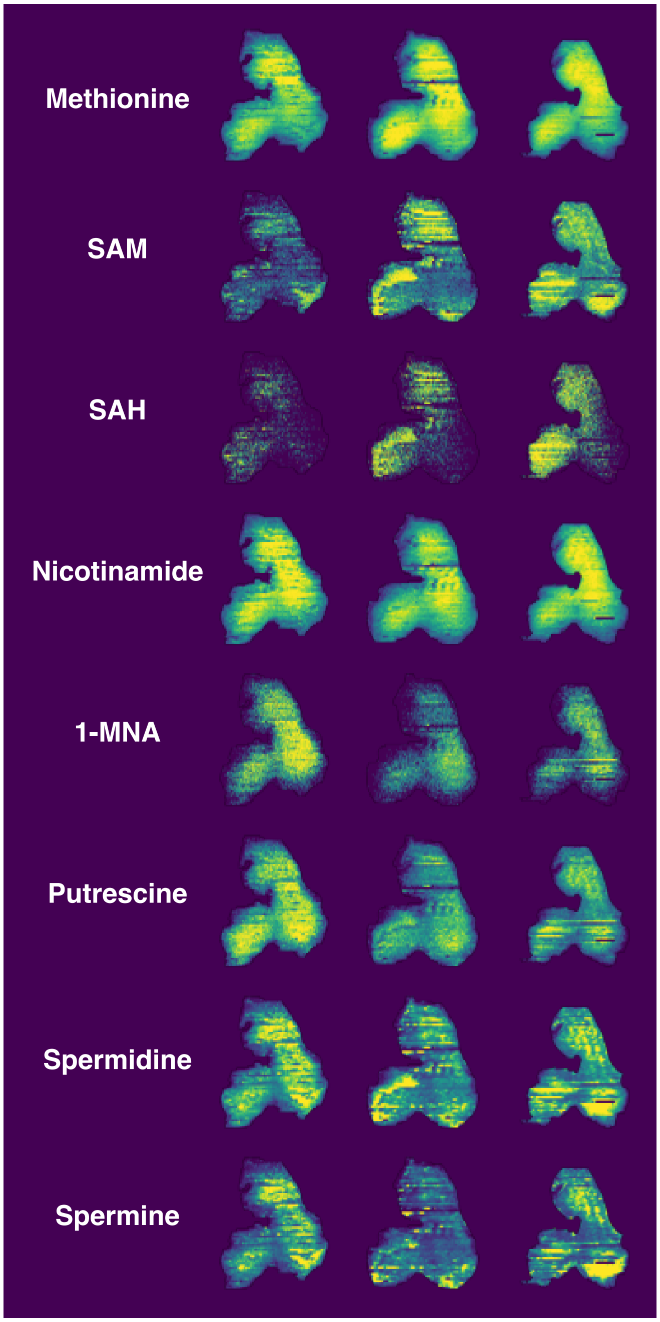


**Figure S11.** Technical reproducibility across three replicates for methionine cycle intermediates in OCT-embedded tissue. A uniform color scale (per metabolite) is applied across the three datasets between the 0–90$^{th}$ percentile. Images are TIC-normalized with a 10ppm tolerance, and an ROI mask was applied to only show tissue-associated pixels. Replicate #3 (3$^{rd}$ column), which is shown in Figure 4 of the main manuscript, was rotated 180° from the original imaging direction for easier comparison between replicates.

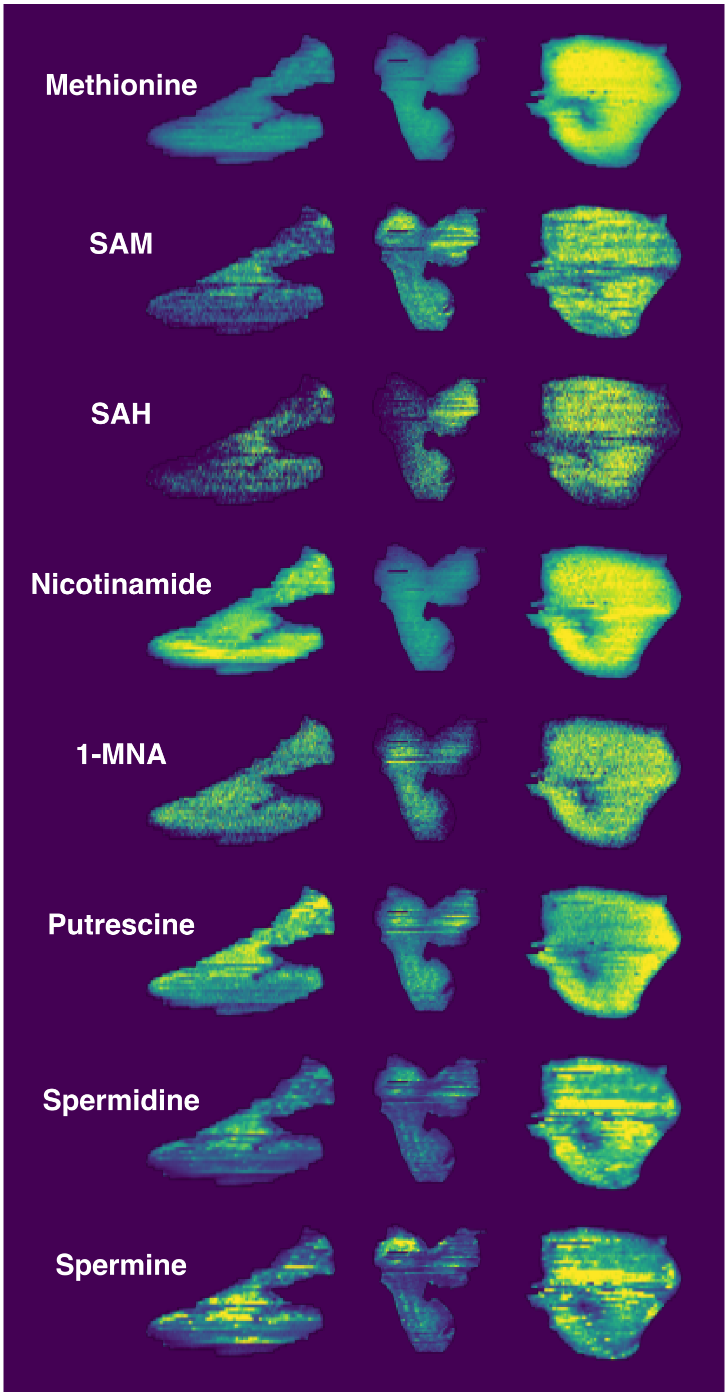


**Figure S12.** Biological variability across three replicates for methionine cycle intermediates in OCT-embedded tissue. A uniform color scale (per metabolite) is applied across the three datasets between the 0–90th percentile. Images are TIC-normalized and generated with a 10ppm tolerance window. An ROI mask was applied to only show tissue-associated pixels.

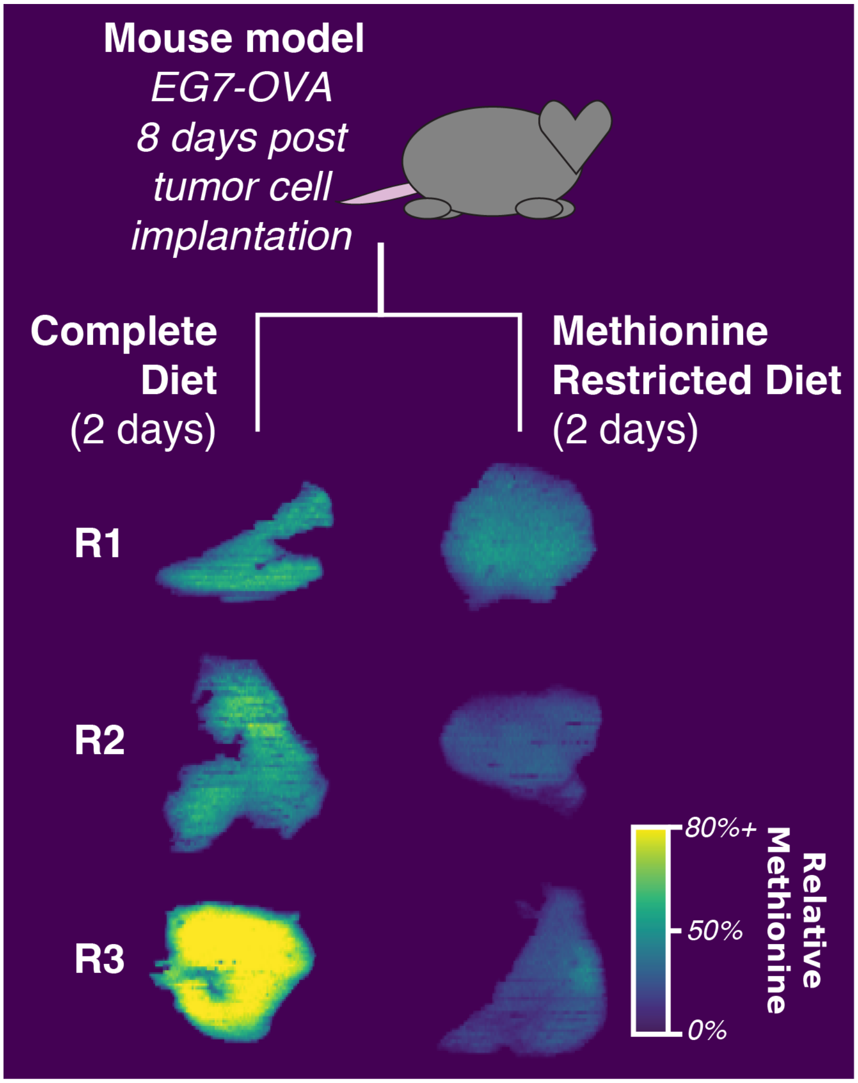


**Figure S13.** Overview of imaging data for methionine in methionine-restricted (MR) and control tumors for OCT embedding. Images are TIC normalized and generated with a 5ppm tolerance window. Color scale is uniform across each embedding material to facilitate easy comparison between control and MR data. CMC images are available in Zhao *et al.* 2025.[5]

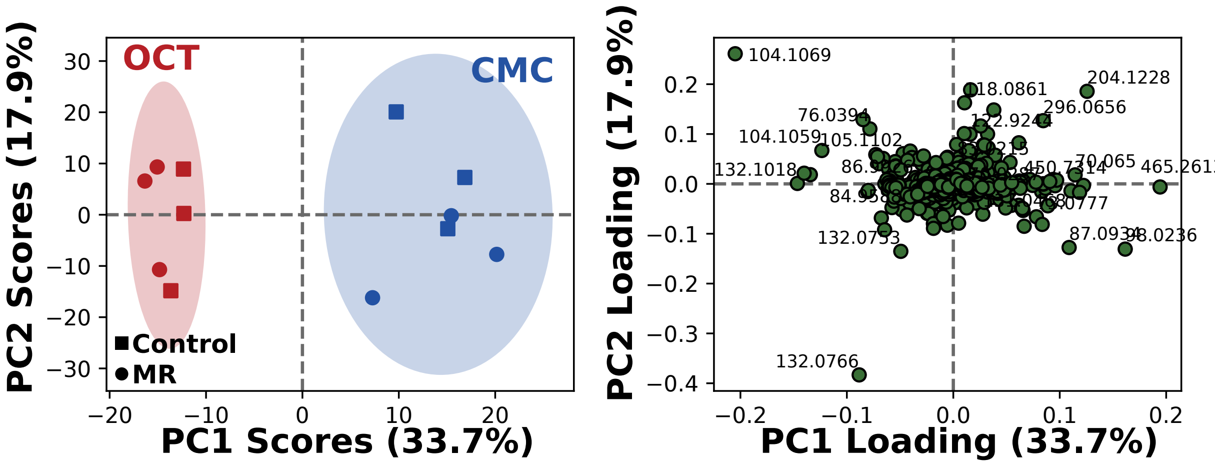


**Figure S14.** PCA Scores and Loadings plot for the average spectra of all the tumors imaged. Data were mean-centred and pareto-scaled prior to PCA using MetaboAnalyst. Markers in red indicate images from OCT-embedded tumors, and those in blue indicate images from CMC-embedded tumors. Square markers show tumors derived from mice fed a control diet, and circle markers represent tumors from mice fed a methionine-restricted diet.

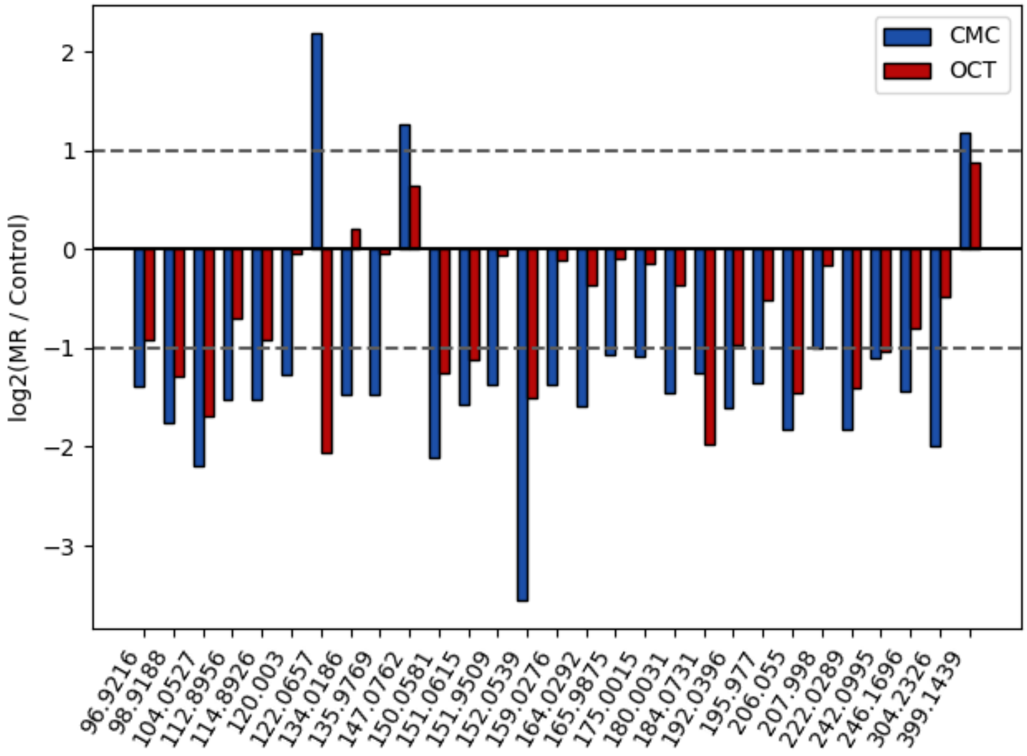


**Figure S15.** Significantly differing features from the CMC-embedded comparison between methionine-restricted (MR) tumors and control compared to the corresponding OCT data. In general, features significant in the CMC show the same direction of change in the OCT-embedded tissue, but do not always reach the threshold for statistical significance ($p = 0.05$; $\log_2$(MR / Control) > 1 or < -1) with the limited sample set in this pilot study ($n = 3$).

**Table S1.** Summary of LC parameters for metabolite annotation in the rat brain and tumor.

| Parameter | Value |
|---|---|
| LC Model | Thermo Fisher Vanquish HPLC |
| Column model | Hypersil GOLD HILIC column (1 x 150 mm) |
| injection volume (μL) | 2 |
| mobile phase A | 20 mM ammonium formate in HPLC-grade water |
| mobile phase B | Acetonitrile |
| Gradient | 1. 0 min, 10% A<br>2. 24 min, 80% A<br>3. 30 min, 10% A |
| post gradient equilibrium | 15 mins at 10% A |
| Ramp | Linear |
| Flow rate | 0.15 mL/min |

**Table S2.** Summary of MS scan parameters for metabolite annotation in the rat brain and tumor.

| Scan Type | Parameter | Value |
|---|---|---|
| Fullscan | Mass window (*m/z*) | 60-900 |
| | Target Resolution (m/Δm @ *m/z* 200) | 120 000 |
| $MS^2$ (targeted and DDA) | isolation window (*m/z*) | 0.5 |
| | Normalized Collision Energy (%) | 15-45 |
| | Target Resolution (m/Δm @ *m/z* 200) | 15 000 |

*Global instrument parameters include capillary voltage: 3.5 kV (+ ESI), -2.5 kV (- ESI); sheath gas: 35 AU, auxiliary gas: 7 AU, ion transfer tube temperature: 320 °C, vaporizer temperature: 275 °C.

**Table S3.** *m/z* peaks in the 480–1000 range that are significantly elevated in the OCT or CMC homogenates in Figure 1.

| m/z | log2(oct / cmc) | OCT crosstalk?* | m/z | log2(oct / cmc) | OCT crosstalk? |
|---|---|---|---|---|---|
| 485.2291 | 2.98 | TRUE | 570.3399 | -1.07 | TRUE |
| 485.5638 | 3.35 | TRUE | 579.3719 | -1.18 | FALSE |
| 487.2524 | 2.93 | TRUE | 582.3084 | -1.10 | FALSE |
| 488.5874 | -1.25 | FALSE | 582.6327 | 2.30 | TRUE |
| 488.9283 | -1.05 | FALSE | 582.6387 | 2.21 | TRUE |
| 489.2764 | 1.61 | TRUE | 583.3101 | -1.45 | FALSE |
| 493.3288 | -1.06 | FALSE | 609.3810 | -1.03 | TRUE |
| 494.5800 | 3.37 | TRUE | 612.3222 | 1.16 | TRUE |
| 495.3450 | -1.13 | FALSE | 612.3450 | -2.06 | FALSE |
| 499.9048 | 3.38 | TRUE | 617.1808 | -1.16 | FALSE |
| 500.2390 | 2.72 | TRUE | 620.3099 | 1.26 | TRUE |
| 502.2930 | -1.05 | FALSE | 620.8116 | 1.31 | TRUE |
| 505.3494 | -1.51 | FALSE | 621.4178 | -1.05 | FALSE |
| 507.3300 | -1.09 | FALSE | 641.3678 | -1.12 | FALSE |
| 509.2553 | 3.32 | TRUE | 642.3229 | 1.67 | TRUE |
| 509.3240 | -1.22 | FALSE | 642.8249 | 1.80 | TRUE |
| 510.3073 | -1.37 | TRUE | 647.3701 | -1.06 | FALSE |
| 514.5802 | 3.61 | TRUE | 664.3364 | 2.09 | TRUE |
| 519.2733 | 1.12 | TRUE | 667.4243 | -1.18 | FALSE |
| 521.3299 | -1.16 | FALSE | 686.3494 | 2.51 | TRUE |
| 523.9309 | 3.41 | TRUE | 703.5476 | -1.09 | FALSE |
| 523.9376 | 2.66 | FALSE | 708.3626 | 2.91 | TRUE |
| 529.2556 | 3.51 | TRUE | 711.4507 | -1.16 | FALSE |
| 533.2822 | 1.09 | TRUE | 722.3984 | -1.02 | TRUE |
| 535.3458 | -1.06 | FALSE | 725.5316 | -1.02 | FALSE |
| 538.6064 | 3.12 | TRUE | 730.5762 | -1.01 | FALSE |
| 538.6129 | 2.69 | FALSE | 731.5790 | -1.28 | FALSE |
| 539.2837 | -1.28 | FALSE | 754.5369 | -1.31 | FALSE |
| 539.9679 | -1.14 | FALSE | 756.5905 | -1.26 | FALSE |
| 543.9311 | 4.15 | TRUE | 780.5909 | -1.02 | FALSE |
| 544.2653 | 3.10 | TRUE | 797.5893 | -1.00 | FALSE |
| 551.3561 | -1.07 | FALSE | 817.6521 | -1.08 | FALSE |
| 553.2834 | 2.73 | TRUE | 831.5727 | -1.00 | FALSE |
| 559.3174 | -1.03 | FALSE | 842.6642 | -1.02 | FALSE |
| 567.9573 | 2.51 | TRUE | 870.6956 | -1.24 | FALSE |
| 567.9635 | 2.41 | FALSE | -- | -- | -- |

*Within 5ppm of a peak observed in the OCT spectra (as M+H, M+Na, or M+K).

**Table S4.** Summary of methionine-cycle intermediates compiled from HMDB.[1]

| Metabolite | HMDB ID | MW (g/mol) | logP | Typical concentration |
|---|---|---|---|---|
| Methionine | HMDB0000696 | 149.211 | -1.8 | 20–30 μM (blood) |
| SAM | HMDB0001185 | 399.445 | -5.3 | 0.05–0.25 μM (blood) |
| SAH | HMDB0000939 | 384.411 | -2.4 | 0.01–0.46 μM (blood) |
| Homocysteine | HMDB0000742 | 135.185 | -2.3 | 8–18 μM (blood) |
| Nicotinamide | HMDB0001406 | 122.123 | -0.4 | 0.03–0.44 μM (blood) |
| 1-MNA | HMDB0000699 | 137.159 | -3.7 | 0.007–0.850 μM (blood) |
| Putrescine | HMDB0001414 | 88.152 | -0.7 | 0.20–0.26 μM (blood) |
| Spermidine | HMDB0001257 | 145.246 | -0.6 | 8.5–10 μM (blood) |
| Spermine | HMDB0001256 | 202.3402 | -0.7 | 0.1–10 μM (blood) |

**Table S5.** Significantly differing features from the CMC-embedded comparison between MR and control and the corresponding data from OCT-embedded analogs.

| | CMC-embedded | | | | OCT-embedded | | | |
|---|---|---|---|---|---|---|---|---|
| *m/z* | **Control Signal** | **MR Signal** | **$log_2$(MR/control)** | **-log(p)** | **Control Signal** | **MR Signal** | **$log_2$(MR/control)** | **-log(p)** |
| 96.9216 | 3.7 | 1.4 | -1.39 | 1.4 | 1.6 | 0.9 | -0.9 | 0.5 |
| 98.9188 | 1.2 | 0.3 | -1.76 | 1.4 | 0.4 | 0.2 | -1.3 | 0.3 |
| 104.0527 | 8.2 | 1.8 | -2.19 | 1.6 | 4.7 | 1.5 | -1.7 | 1.4 |
| 112.8956 | 12.7 | 4.4 | -1.52 | 1.3 | 4.0 | 2.4 | -0.7 | 0.4 |
| 114.8926 | 3.9 | 1.4 | -1.52 | 1.3 | 1.2 | 0.6 | -0.9 | 0.4 |
| 120.0030 | 1.7 | 0.7 | -1.27 | 2.0 | 0.3 | 0.3 | 0.0 | 0.0 |
| 122.0657 | 0.1 | 0.4 | 2.19 | 1.5 | 0.4 | 0.1 | -2.1 | 0.6 |
| 134.0186 | 1.2 | 0.4 | -1.47 | 1.4 | 0.3 | 0.3 | 0.2 | 0.1 |
| 135.9769 | 3.7 | 1.3 | -1.47 | 1.7 | 1.3 | 1.3 | 0.0 | 0.0 |
| 147.0762 | 6.7 | 16.1 | 1.26 | 1.4 | 9.3 | 14.5 | 0.6 | 0.3 |
| 150.0581 | 32.5 | 7.6 | -2.10 | 4.1 | 37.7 | 15.8 | -1.3 | 1.3 |
| 151.0615 | 1.8 | 0.6 | -1.58 | 3.5 | 2.2 | 1.0 | -1.1 | 1.6 |
| 151.9509 | 4.2 | 1.6 | -1.37 | 1.3 | 1.4 | 1.3 | -0.1 | 0.0 |
| 152.0539 | 1.3 | 0.1 | -3.55 | 3.2 | 1.5 | 0.5 | -1.5 | 1.3 |
| 159.0276 | 3.0 | 1.1 | -1.38 | 2.0 | 3.7 | 3.4 | -0.1 | 0.1 |
| 164.0292 | 1.8 | 0.6 | -1.59 | 1.5 | 0.2 | 0.2 | -0.4 | 0.1 |
| 165.9875 | 2.0 | 1.0 | -1.06 | 1.4 | 0.8 | 0.8 | -0.1 | 0.1 |
| 175.0015 | 7.4 | 3.5 | -1.08 | 1.7 | 10.0 | 9.0 | -0.1 | 0.1 |
| 180.0031 | 4.5 | 1.7 | -1.45 | 2.1 | 1.5 | 1.2 | -0.4 | 0.2 |
| 184.0731 | 12.5 | 5.2 | -1.26 | 1.8 | 14.2 | 3.6 | -2.0 | 2.1 |
| 192.0396 | 0.9 | 0.3 | -1.61 | 1.7 | 0.1 | 0.1 | -1.0 | 0.2 |
| 195.9770 | 3.0 | 1.2 | -1.36 | 1.5 | 0.9 | 0.7 | -0.5 | 0.2 |
| 206.0550 | 10.8 | 3.0 | -1.83 | 1.4 | 5.0 | 1.8 | -1.5 | 2.5 |
| 207.9980 | 4.4 | 2.2 | -1.01 | 2.1 | 2.1 | 1.9 | -0.2 | 0.1 |
| 222.0289 | 17.3 | 4.9 | -1.83 | 1.3 | 8.7 | 3.3 | -1.4 | 1.2 |
| 242.0995 | 1.3 | 0.6 | -1.10 | 2.2 | 0.4 | 0.2 | -1.0 | 0.4 |
| 246.1696 | 1.8 | 0.7 | -1.44 | 1.5 | 0.7 | 0.4 | -0.8 | 0.4 |
| 304.2326 | 0.7 | 0.2 | -1.99 | 1.4 | 0.6 | 0.4 | -0.5 | 0.2 |
| 399.1439 | 1.2 | 2.8 | 1.18 | 2.4 | 1.4 | 2.5 | 0.9 | 1.0 |
